# An individual interneuron participates in many kinds of inhibition and innervates much of the mouse visual thalamus

**DOI:** 10.1101/683276

**Authors:** Josh L. Morgan, Jeff W. Lichtman

**Affiliations:** John F. Hardesty, MD Department of Ophthalmology and Visual Science and Department of Neuroscience. Department of Biomedical Engineering. Washington University School of Medicine, St. Louis MO 63108; Department of Molecular and Cell Biology and Center for Brain Science, Harvard University, Cambridge MA 02138

## Abstract

One way to assess a neuron’s function is to describe all its inputs and outputs. With this goal in mind, we used serial section electron microscopy to map 899 synaptic inputs and 623 outputs in one inhibitory interneuron in a large volume of the mouse visual thalamus. This neuron innervated 256 thalamocortical cells spread across functionally distinct subregions of the visual thalamus. All but one of its neurites were bifunctional, innervating thalamocortical and local interneurons while also receiving synapses from the retina. We observed a wide variety of local synaptic motifs. While this neuron innervated many cells weakly, with single *en passant* synapses, it also deployed specialized branches that climbed along other dendrites to form strong multi-synaptic connections with a subset of partners. This neuron’s diverse range of synaptic relationships allows it to participate in a mix of global and local processing but defies assigning it a single circuit function.

## INTRODUCTION

A legacy of Ramón y Cajal’s analysis of Golgi stained tissue is the Neuron Doctrine stating that the nervous system is made up of discrete cells that act as the functional units of the brain. Critical to this unitary function of neurons is the law of dynamic polarization that states that information flows through neurons from dendrites to soma to axon. This framework, however, breaks down when considering neurons where inputs and outputs intermingle on the same neurites. In these cases, individual neurites of the same cell can process and transmit different signals by virtue of their distinct input and output connectivities. Neurons with intermingled inputs and outputs include amacrine cells of the vertebrate retina (Grimes et al., 2010; Hausselt et al., 2007; Kidd, 1962), thalamic interneurons (Carden and Bickford, 2002; Famiglietti, 1970; Morest, 1975), granule cells of the olfactory bulb (Rall et al., 1966) and many invertebrate neurons (Christiansen et al., 2011; Graubard and Calvin; Gray, 1969; White et al., 1986). In addition, many mammalian cells with clear input neurites (i.e. dendrites) and output neurites (i.e. axons) receive some degree of axo-axonic (Strettoi et al., 1990) or dendro-dendritic (Hattori et al., 1979; Sloper and Powell, 1978) innervation. To get a better idea of how to describe the circuitry of a neuron with pre- and postsynaptic mixing on its neurites we sought to identify all the presynaptic input and all the postsynaptic output of one thalamic local interneuron (LIN) known to have mixing based both on anatomical (Famiglietti, 1970; Famiglietti and Peters, 1972; Guillery, 1972; Güillery, 1969; Rafols and Valverde, 1973) and physiological (Blitz and Regehr, 2005; Cox and Sherman, 2000; Crandall and Cox, 2012, 2013; Govindaiah and Cox, 2004) studies.

Here, by using serial electron microscopy to trace out almost all of a LIN’s neurites and identify all the associated pre- and postsynaptic sites, we sought to obtain a complete picture of the input/output connections of a single neuron. This analysis revealed that the LIN participated in hundreds of different neuronal interactions using many kinds of synaptic motifs that spanned many functional domains in the visual thalamus.

## RESULTS

### One LIN projects throughout the dLGN

We previously reconstructed the retinal ganglion cell (RGC) to thalamocortical cell (TC) connectivity in a volume of the postnatal day 32 mouse dLGN (Morgan et al., 2016). While additional synaptic rearrangement occurs in the second month of dLGN postnatal development (Hong et al., 2014), P32 is a post weanling juvenile that can forage for food independently. For this study, we used this same “digital tissue” (Morgan and Lichtman, 2017). Segmentations were performed on an aligned image volume downsampled to 16 × 16 x 30 nm (Full resolution available at http://bossdb.org/project/morgan2020). Because the acquired volume is in the temporal region of the dLGN, it is innervated monocularly by RGCs from the contralateral eye (Jaubert-Miazza et al., 2005). In the dLGN, there are two types of neuronal somata: excitatory TCs and inhibitory LINs. LINs are easily differentiable from the TCs by their geometry (no myelinated axon, smaller cell body, non-tapering neurites)(Zhu and Lo, 1999), connectivity and ultrastructure (Sherman, 2004). We chose one LIN for detailed analysis (LIN1). LIN1 was chosen because it’s cell body is near the middle of 10,000 30 nm sections image volume (400 μm x 600 μm x 300 μm) and it is connected by synapses to the previously reconstructed network of RGC axons and TCs (Morgan et al., 2016).

LIN1 spanned the entire depth of the dLGN (500 μm) and nearly the entire breadth of the image volume (400 μm, exiting the lateral and medial borders) and had a total reconstructed neurite length of 5999 μm. This large arborization is consistent with previous descriptions of mouse (Seabrook et al., 2013) and rat LINs (Williams et al., 1996; Zhu and Lo, 1999; Zhu and Uhlrich, 1997; Zhu et al., 1999). Given the ∼1 mm width of the dLGN in this 32 day-old mouse, this arbor expanse covers at least half the visual field (Figure 1). The vertical extent of this arborization means that its arbor also overlaps multiple channels of retinal input and, notably, overlaps both the core and shell regions of the dLGN. These regions delineate the important functional division between the recipient zones for direction selective RGCs (shell) and primarily non-direction selective RGCs (core) in mouse dLGN (Reese, 1988). If all these neurites are synaptically connected to nearby neurons (see below), then LIN1 is situated so that it has the potential to influence circuits with different functions. A second tracer performed an independent partial reconstruction of LIN1’s arbor and found a matching widespread distribution of neurites (Supplemental Figure 1A). The wide arborization of this LIN does not appear to be exceptional; the other LINs that we reconstructed also spanned large areas of the dLGN (Figure 1B; Supplementary Movie 1).

**Figure 1:**
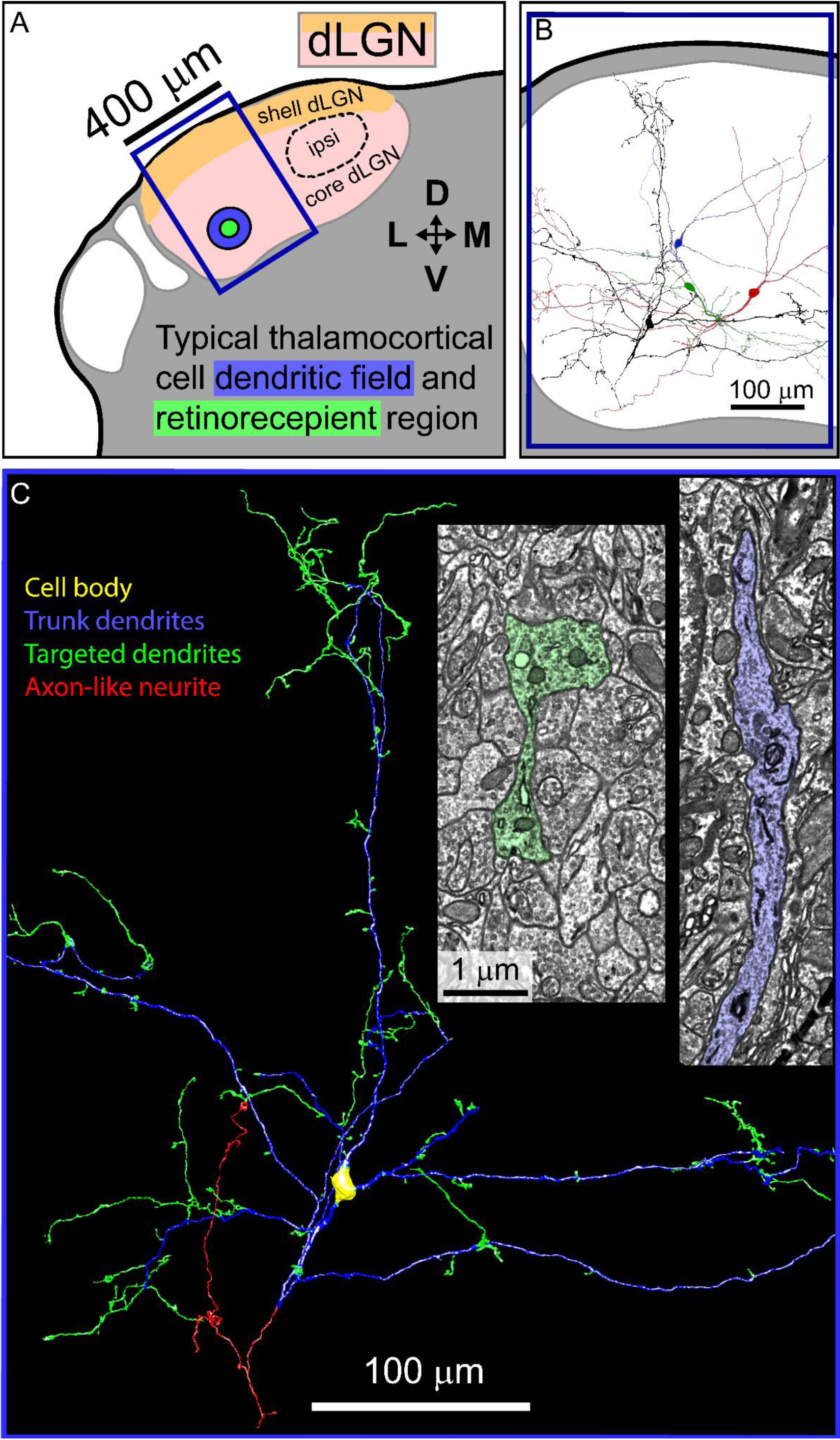
Morphology of dLGN LINs. A) EM volume (blue box) relative to a coronal view of dLGN. Blue box indicates the EM volume and is the same area as the blue box in B and C. Typical size of TC dendritic arbor (blue) and the proximal retinorecepient region of a TC (green) relative to the dLGN (core = pink, shell = orange, dotted line = ipsilateral). B) Partial tracings of five LINs. LIN1 is black. C) LIN1 from panel C: yellow cell body, blue trunk dendrites, green targeting dendrites, red axon-like neurite. Insets = electron micrographs of trunk (blue) and targeting (green) dendrites.

The neurites of LIN1 were divided into three distinct types (Figure 1C) which appeared homologous to the trunk dendrites, dendritic appendages and axon described previously in cats (Hamos et al., 1985; Montero, 1991). Here we identified *trunk dendrites* as the relatively thick and straight neurites (mean width 0.93 ±0.13 μm, 1107 measures, tortuosity 1.048 ± 0.001, 627 measures) that extend for hundreds of micrometers and that traverse most of the span of LIN1’s territory. We used the convention *dendrite* to refer to these processes but note that the trunk dendrites form more output synapses than input synapses (see below). The three thick primary neurites emerging from LIN1’s cell body were trunk dendrites. In contrast, we defined *targeting dendrites* as the significantly thinner more circuitous neurites (mean width 0.64 ± 0.01 μm, 1341 measures, P < 0.001, tortuosity 1.211 ±0.018, n = 490 measures, P < 0.001, Supplementary Figure 1B) with multiple irregularly shaped swellings interlinked by thinner neurites. The targeting dendrites ranged from short, single bouton spine-like neurites to long (>50 um) branched structures. Targeting dendrites emerged from all regions of the trunk arborization. Trunk dendrites were not observed emerging from targeting dendrites. The axon, identified primarily by its lack of RGC inputs, resembles small LIN axons previously reported in rats (Zhu and Lo, 1999) and cats (Montero, 1987). It consisted of a relatively small arbor (366 μm) with five terminal neurites and a small side filopodium (Figure 1C). The axon emerged from a tertiary trunk dendrite 67.4 μm away from the cell body and it terminated within the imaged volume. Morphologically, the axon was varicose and thin (0.45 ±0.02 μm) and resembled the targeting dendrites of LIN1. Analysis of LIN1’s synaptic connections (below) revealed that the three types of neurites – axon, trunk dendrite, and targeting dendrite-represent three modes of innervating the dLGN network.

### One LIN forms pre- and postsynaptic connections throughout the dLGN

To determine if the expansive LIN arbor was in fact linked by synapses to multiple regions of the dLGN, we mapped all LIN1’s synapses contained in the 100 TB volume (Supplementary Figure 2A, Supplementary Movie 2). We identified synapses as sites where LIN1 was in close apposition to another cell and where ∼40 nm diameter vesicles (putative synaptic vesicles) were clustered against one of the apposed membranes (Figure 2). To confirm that our use of downsampled data did not miss synapses, we compared synapse identification within full resolution data (8 × 8 x 3 μm volume at 4 × 4 x 30 nm resolution) to a down-sampled version of the same volume (at 16 × 16 x 30 nm resolution, Supplementary Figure 3). We found 94.6% of 168 synapses identified in the high-resolution dataset were also identified in the down-sampled dataset. To control for specificity, we rotated the downsampled segmentation by 90 degrees; only 19.4% of high-resolution synapses overlapped with the low-resolution segmentation.

**Figure 2.**
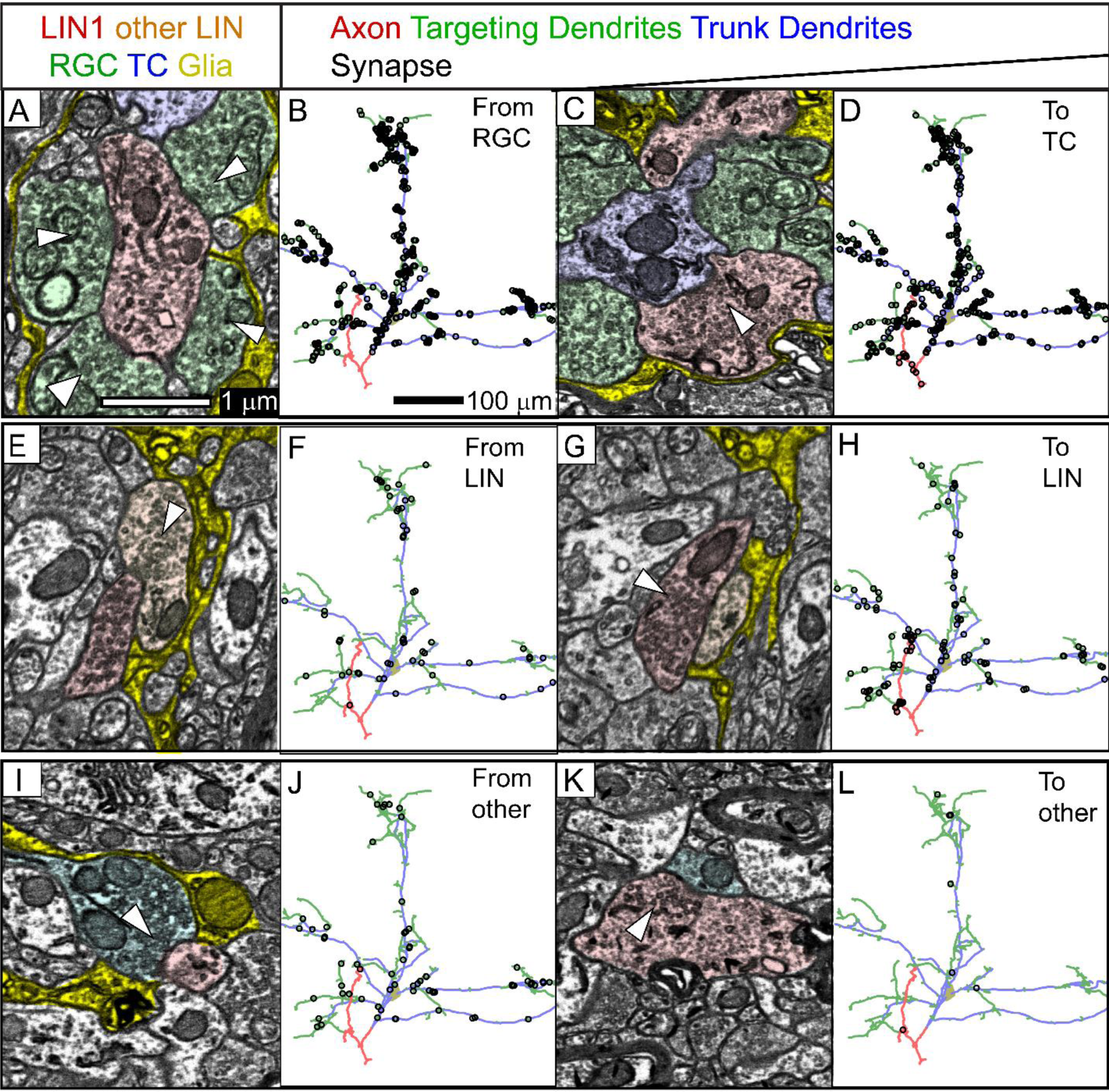
LIN1 forms pre and postsynaptic contacts throughout its arbor. In EM: presynaptic vesicles clusters = white arrows, Glial sheaths = yellow, TCs = blue, LIN1 = red, other LINs = orange, RGCs = green, and unidentified types = cyan. A) EM showing RGC input to LIN1. B) Distribution of RGC inputs to LIN1. C) EM showing LIN1 intputs to TC. D) Distribution of LIN1 inputs to TCs. E) EM showing LIN1 input to second LIN. Synapse in E and G are reciprocal synapses separated by 330 nm. F) Distribution of LIN1 inputs to LINs. G) LIN input to LIN1. H) Distribution of LIN inputs to LIN1. I) EM showing unidentified inputs to LIN1. J) Distribution of unidentified inputs to LIN1 synapses. K) LIN1 input onto an unidentified cell type. L) Distribution of LIN1 inputs to unidentified cell types.

In terms of input, we identified multiple sources that established synapses with this cell. There were 775 sites where RGC axons established synapses onto LIN1, 70 synapses onto LIN1 from other LINs, and 54 synapses for which the input cell type could not be definitively identified (Figure 2, Table 1). The RGC inputs included synapses from both large and small bouton forming RGCs (Morgan et al., 2016). Thirty-six of LIN1’s connections to other LINs were reciprocal synapses and four were autaptic synapses that LIN1 formed with itself. Our measure of 70 LIN synapses presynaptic to LIN1 is likely an undercount because of their relatively small and relatively undifferentiated morphology (Figure 2E, G). In addition to RGC and LIN inputs, LINs are known to receive input from parabranchial brainstem axons (McCormick and Pape, 1988), pretectogeniculate neurons (Wang Bickford 2002), corticothalamic inputs (Weber et al., 1989) and other cell types. Here, we group all these inputs together as ‘unidentified’. However, the most common unidentified inputs were ultrastructurally consistent with cholinergic brainstem axons, possessing large oval presynaptic boutons with dark cytosol (Erişir et al., 1997). We traced 14 of these unidentified axons back to synapses they formed on TCs and found that 12 formed boutons that resembled the synapses they formed on LIN1. Two formed small boutons on TC shafts consistent with thalamic reticular nucleus inputs. None formed small boutons on distal TC spines (consistent with cortical input (Guillery, 1971). LIN1, therefore, did not appear to receive much, if any, of the excitatory cortical feedback that dominates the distal dendrites of TCs. In terms of output, LIN1 established 492 synapses presynaptic to TCs, 124 synapses presynaptic to LINs (including the four onto itself), and 7 synapses presynaptic to unidentified cell types (Figure 2). There was an average of one incoming RGC synapse per 7.7 μm of the LIN neurite length and one outgoing TC synapse per 12.2 μm of neurite length, although these numbers belie the fact that the synapses were unevenly distributed (see below).

**Table 1.** Summary of synaptic contacts for LIN1 divided by whole arbor, trunk dendrites, targeting dendrites, and the axon-like neurite.

|  | Full cell | Trunk Dendrites | Targeting Dendrites | Axon |
| --- | --- | --- | --- | --- |
| Average cross section area | 43.8 $\mu\text{m}^2$<br>(cell body) | 0.93 $\mu\text{m}$ | 0.64 $\mu\text{m}$ | 0.45 $\mu\text{m}$ |
| Total length | 5999 $\mu\text{m}$ | 2493 $\mu\text{m}$ | 2784 $\mu\text{m}$ | 366 $\mu\text{m}$ |
| input synapses (per mm) | 899 (150) | 163 (65) | 730 (262) | 2 (5) |
| output synapses (per mm) | 623 (104) | 201(81) | 378 (136) | 44 (120) |
| RGC inputs (per mm) | 775 (129) | 129 (52) | 642 (231) | 0 |
| RGC inputs with TC outputs within 5 $\mu\text{m}$ (per mm) | 654 (109) | 117 (47) | 537 (193) | 0 |
| TC outputs (per mm) | 492 (82) | 171 (69) | 299 (107) | 22 (60) |
| TC outputs with RGC inputs within 5 $\mu\text{m}$ (per mm) | 385 (64) | 113 (45) | 272 (98) | 0 |
| Thalamocortical cells identified as targets (per mm) | 256 (43) | 123 (49) | 146 (52) | 13 (36) |
| LIN inputs (per mm) | 70 (12) | 12 (5) | 56 (20) | 1 (.4) |
| LIN outputs (per mm) | 124 (20) | 27 (11) | 76 (27) | 21 (57) |
| Reciprocal LIN->LIN synapses (per mm) | 36 (6) | 7 (3) | 27 (10) | 2 (5) |
| Autaptic synapses (per mm) | 4 (0.6) | 1 (.4) | 3 (1) | 0 |
| Other inputs (per mm) | 54 (9) | 21 (8) | 32 (12) | 1 (.4) |
| Other outputs (per mm) | 7 (1) | 3 (1) | 3 (1) | 1 (.4) |

While non-synaptic adherins junctions are common on TC cells (Morgan et al., 2016), we did not observe these sorts of junctions on LIN1.

Although no part of LIN1 was myelinated, most of its surface was ensheathed by glia. This wrapping was continuous with the glial ensheathment covering the synaptic glomeruli innervated by LIN1 (Supplementary Figure 2E). Although the glial ensheathment was shared with RGC boutons innervating the LIN, the ensheathment did not extend to include TC dendrites innervated by LIN1. At sites where LIN1’s trunk dendrites formed and received synapses, the glial ensheathment typically extended to the edges of the synaptic apposition leaving only a small gap in the glial sheath (Supplementary Figure 2F). Whether the glial ensheathment is there to reuptake GABA (De Biasi et al., 1998; Errington et al., 2011), shape membrane capacitance, or perform some other function remains to be determined, but for whatever reason, this glial association is strikingly different from the way glial processes and TC dendrites interact.

Every major neurite of LIN1, both trunk and targeting, innervated TCs and thus all of LIN1’s neurites have axon-like properties: 94.3% of the arbor was within 20 μm of a site presynaptic to a TC dendrite. Although LIN1 provided presynaptic input to TC dendrites throughout its widely distributed arbor, it remained possible that the dendrites being innervated were derived from a small subset of TC cell bodies that were more restricted in location or type. To determine the number and distribution of TC somata innervated by LIN1, we traced the TC dendrites postsynaptic to LIN1->TC synapses back to their cell bodies. We found 256 TCs innervated by LIN1. These neurons were distributed widely throughout the thalamus. Typically, the cell bodies were located near the sites of LIN1’s inhibitory contact (median distance between a TC soma and a LIN1 synapse = 46 μm, 95% range 17-122 μm, n = 394 synapses, Supplemental Figure 2G). The resulting spatial distribution of TCs innervated by the LIN, therefore, roughly followed the distribution of the LIN arbor itself (Figure 3A).

**Figure 3.**
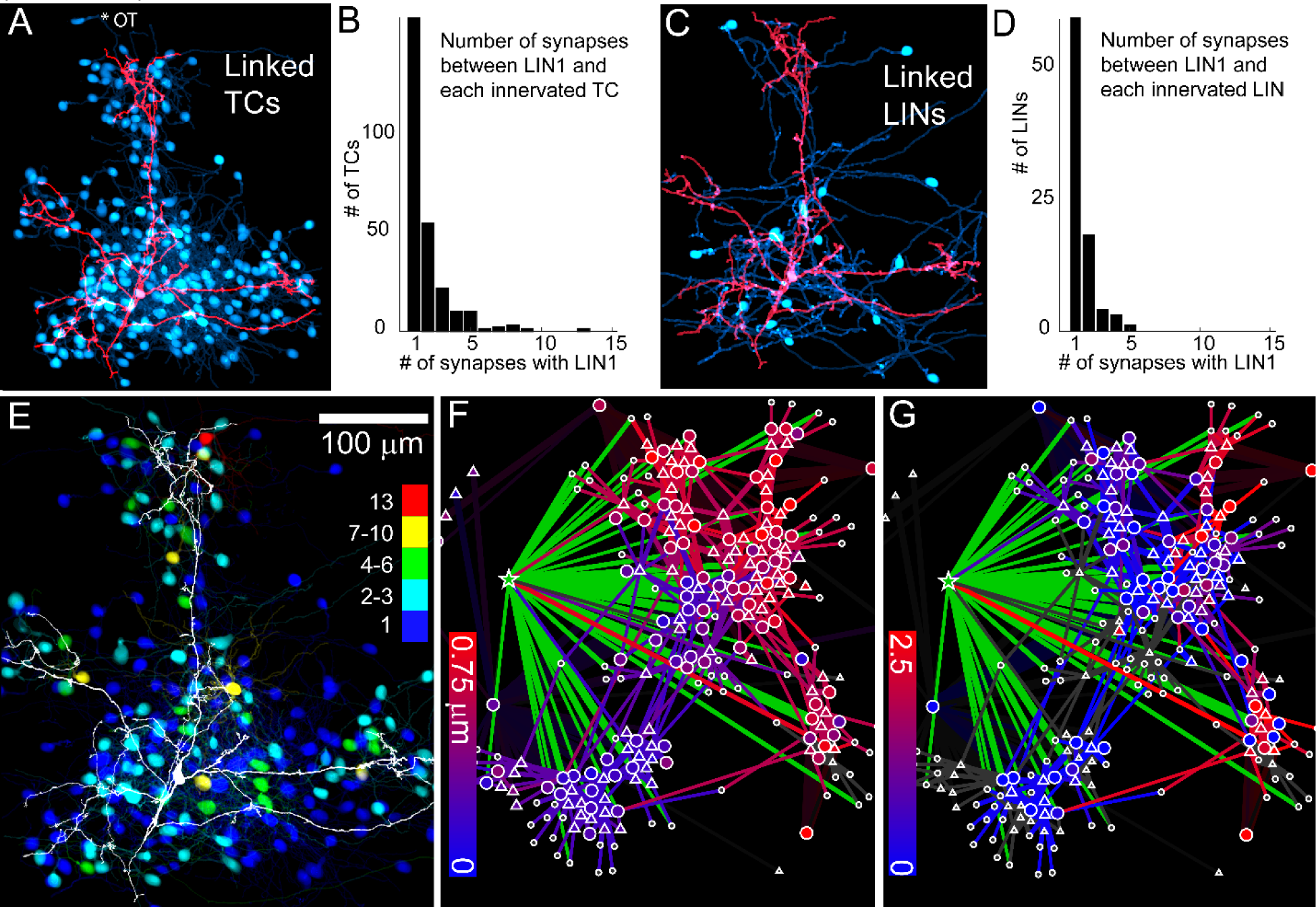
LIN1 connects to partners throughout the dLGN. A) Distribution of cell bodies (cyan) of TCs that are innervated by LIN1 (red). B) Histogram showing number of synapses connecting LIN1 to TC partners. C) Distribution of cell bodies of LINs that could be traced from their synapses with LIN1. D) Histogram showing number of synapses connecting LIN1 to LIN partners. E) Spatial distribution of TCs that receive many (red, yellow) or few (blue, cyan) synapses from LIN1. F) Links of LIN1 (green star) to all clusters within a force-directed sorting of a synaptically connected network of RGCs (triangles) and TCs (circles). RGC bouton diameter associated with RGCs and TCs is coded from small (blue) to large (red). Thicker lines represent more synapses between cell pairs. G) Links of LIN1 (green star) to all clusters within network from panel F. TC spine perforation of RGC boutons is coded from none (blue) to several (red).

We also found both single synapse and multiple synapse links between LIN1 and individual TCs throughout LIN1’s arbor. Sixty percent (160) of LIN1’s TC targets received only one synapse from LIN1 (Figure 3B). However, a substantial number of LIN1’s associations with TCs were much stronger: 32% of the input to TCs were to TCs receiving 4 or more synapses from LIN1. The TCs that received the strongest input from LIN1 were not concentrated in any region of the dLGN arbor. The 8 TCs that received the largest numbers of synapses (13,9,8,8,8,7,7,6) were distributed roughly evenly across the LIN1 arborization field and are therefore likely to be innervated by different types of RGCs representing different regions of visual space (Figure 3E). The wide expanse of this arbor allowed this single LIN to establish strong connections to TCs at nearly opposite ends of the dLGN in both the mediolateral and dorsoventral axis. Thus, this neuron provided inhibition to TCs that have receptive fields dispersed throughout the visual field of the contralateral eye. Because these TCs were also distributed across the depth of the dLGN they included TCs responding to different channels of visual information (Piscopo et al., 2013; Reese, 1988).

Nearly all neurites of LIN1 also received synaptic input and hence had dendrite-like properties: 86.1% of LIN1’s arbor was within 20 μm of an RGC input. These inputs spanned the full depth and width of its arbor and thus the full depth and most of the width of the dLGN itself. This widespread input distribution is consistent with innervation from RGCs covering much of the visual field and with different receptive field properties.

We identifed one branch of LIN1 that was axon-like in that it was not innervated by RGC axons (Figure 1c). This neurite consisted of several connected branches with a combined length of 366 μm (out of 5999 μm total reconstructed arbor) (Figure 1C). This neurite received one proximal synapse that might have been a cholinergic brainstem input (Erişir et al., 1997) and one LIN input. The lack of RGC innervation on this neurite was not because of spatial inaccessibility. RGC boutons did come into membrane contact with this axon-like neurite and many of these were near LIN1 synapses to TCs. To test the probability of an input-free neurite occurring by chance, we randomized the position of RGC inputs across LIN1’s arbor and counted the number of RGC inputs on the least innervated of 27 axon-sized stretches of neurite. In 10,000 repetitions, we never observed an axon-sized neurite that lacked RGC inputs (average RGC inputs on least innervated neurite = 35.7, 95%CI 29-41, P < 0.001). Ultrastructurally the synapses the axon formed with other LINs and TCs resembled those formed by the targeting neurites elsewhere on LIN1 (Supplementary Figure 2B-D). In terms of output, the axon-like neurite was responsible for only 4% (20 TCs) of LIN1’s identified output to TCs, but a slightly larger proportion (17%; 21 LINs) of LIN1’s output to other LINs. The TCs innervated by this neurite appeared to be functionally heterogeneous because some had small glomerular type RGC input and others had large perforated RGC boutons corresponding to different classes of RGCs and TCs (Morgan et al., 2016). Hence the axon-like process did not appear to restrict its innervation to a single channel of visual processing.

Both pre- (124 synapses) and postsynaptic (70 synapses) connections to other LINs were distributed throughout LIN1’s arbor (Figure 2E-H). The neurites connecting these synapses to the cell bodies of other LINs were much longer than those linking the traced LIN to TCs. Indeed, 69% (38/55) of the LIN dendrites synaptically linked to LIN1 exited the volume before their bodies could be found. The 17 LIN cell bodies that were synaptically connected to LIN1 were 42 μm to 232 μm away (Supplemental Figure 2D) and distributed throughout the entire dLGN volume. Therefore, the LIN->LIN network was significantly more expansive than the LIN->TC network (ranksum synapse to cell body distance for TCs and LINs P< 0.001, Figure 3AC), with LINs having synaptic access to many LINS from other regions of the dLGN. Thus, at the level of cell positioning, mouse LIN-LIN connectivity did not seem constrained by the spatial/functional organization of the visual thalamus.

Despite the wide expanse of the dLGN innervated by LIN1, it was possible that its synapses participated in only specific functional subnetworks. For example, our previous reconstruction of the retinal input to TCs in this same tissue revealed several ultrastructurally distinct subnetworks that were spatially intermixed (Morgan et al., 2016). We therefore testeed whether LIN1 selectively synapsed with a subset of these previously defined subnetworks. The previously reconstructed region included two morphologically distinct groups of retinal axons (large and small bouton-forming) that established synapses onto two morphologically distinct groups of TCs: those that received perforated and those that received exclusively non-perforated RGC synapses. We found that LIN1 innervated all parts of the previously traced RGC to TC network (Figure 3F, G). In particular, LIN1 was synaptically associated with both the large and small bouton forming sets of RGCs (Figure 3F) and both sets of TCs (Figure 3G). These results show that one inhibitory neuron can participate in processing visual information from many distant regions of the visual field and from multiple “parallel” streams of visual processing.

### One LIN generates diverse synaptic motifs

The synaptic glomeruli of the dLGN provide spatially compact structures (<10 um) in which large numbers of synapses might influence each other’s functioning (Famiglietti and Peters, 1972; Guillery and Colonnier, 1970) and many multi-neuronal synaptic motifs have been described for LINs as a class (Cox and Beatty, 2017; Crandall and Cox, 2013; Hámori et al., 1974; Hamos et al., 1985; Hirsch et al., 2015; McCormick and Pape, 1988; Pasik et al., 1976; Sherman, 2004; Weber et al., 1989). These motifs often consist of serial synapses (Colonnier and Guillery, 1964; Hámori et al., 1974), inputs onto LINs that are immediately adjacent to the output synapse from the LIN. Because we had access to more than a thousand synapses to and from this one cell, we could see the degree to which this one neuron participated in one or more kinds of inhibitory synaptic relationships. We found a diverse array of synaptic relationships associated with LIN1 including autaptic synapses, synapses with TCs with no other input nearby, RGC->LIN->TC triadic synapses, LIN->LIN->TC triadic synapses, BS (brainstem)->LIN->TC triad synapses, one-way LIN->LIN synapses and reciprocal LIN<->LIN synapses (Figure 4).

**Figure 4.**
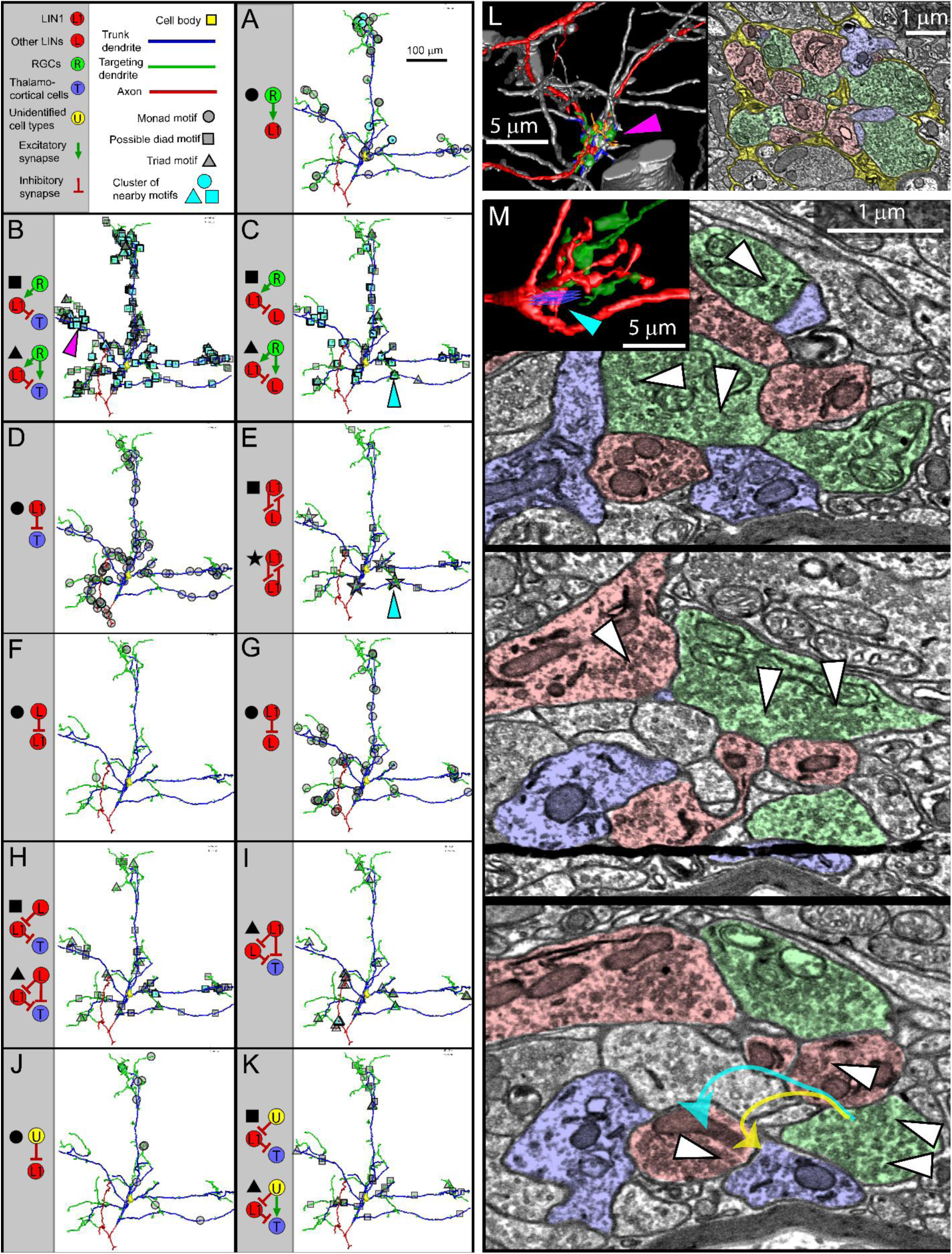
Diverse synaptic motifs of LIN1. (A-J) Motifs identified by searching for combinations of synaptic connections that could be found within 5 μm of one another. (A,F,J) Inputs to LIN1 that were found >5 μm away from TC output synapses. (D) LIN->TC synapses found >5 μm away from an RGC input. A clustered motif (cyan fill) is defined as a motif located <5 μm from at least two examples of the same motif. Magenta and Cyan arrows indicate the position of the synaptic structures shown in L and M respectively. (L) Three-dimensional rendering (left) and EM slice (right) of a synaptic glomerulus of LIN1. Encapsulating glial sheaths are highlighted in yellow, TCs in blue, LIN1 in red, other LINs in orange, RGCs in green and unidentified types in cyan. In 3D rendering, neurites outside of the glomerulus (except for LIN1) are gray. (M) 3D rendering (left) and EM slices (right) showing several synaptic motifs in a LIN1 glomerulus. Blue squares in rendering indicate the position of the EM slices. White triangles indicate synapses. Red profiles are LIN1. Yellow arrow shows path from RGC to LIN1 to TC. Cyan arrow shows path from RGC to LIN1 to LIN1 to TC via an autaptic synapse.

#### RGC->LIN->TC triads motif

The most prevalent kind of inhibitory motif associated with LIN1 was the RGC->LIN->TC triad in which a synaptically connected LIN1 neurite and TC dendrite are both innervated by the same RGC axon, often via the same presynaptic RGC bouton (Figure 4B, 5A, B). This motif has the effect of shortening the duration of the excitation of a TC by a retinal axon by following the retinal excitation with LIN-based inhibition (i.e. temporal sharpening, (Blitz and Regehr, 2005)). RGC->LIN->TC triads could be isolated or clustered together with other synaptic triads (Figure 4B). These clusters of triads were associated with the classic clustered bouton profiles of dLGN glomeruli (Figure 4L, M, Supplementary Movie 3).

To determine the proportion of RGC inputs onto LIN1 that were associated with synaptic triads, we randomly selected 79 of the RGC inputs to LIN1 for additional local circuit analysis. For each of these boutons, we searched the bouton and nearby RGC axon for synapses onto TCs. We found that nearly all (74/79) of the RGC boutons innervating LIN1 served a dual purpose as they also formed synapses with TC dendrites. In cases where RGC input to LIN1 was not associated with a nearby postsynaptic TC neurite (5/79 sampled RGC inputs), the RGC bouton presynaptic to LIN1 was smaller in volume and lacked the high vesicle and mitochondrial densities typical of RGC boutons associated with triads. Of the dual-purpose RGC boutons, most (71/74) formed a synapse onto the TC that was also innervated by LIN1 (hence a triadic motif). In almost all these triadic relationships (69/71), the distance between the RGC input synapse to LIN1 and the TC output synapses from LIN1 was <5 μm (Figure 5D). Therefore, most (84%) of retinal inputs to LIN1 were associated with a compact triadic subcellular pathway of less than 5 μm (Figure 5C, D).

**Figure 5.**
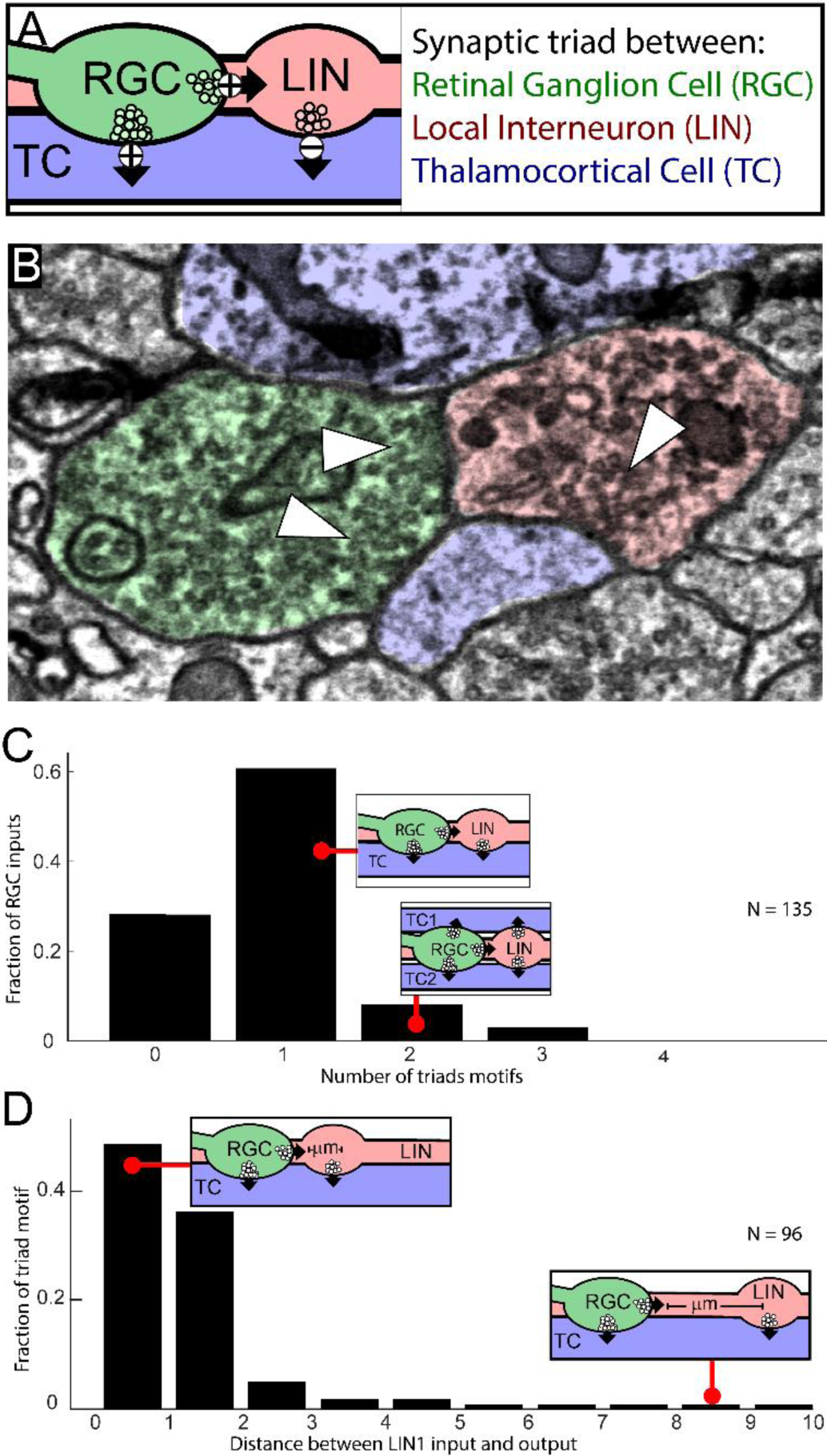
Structure of RGC->LIN->TC synaptic triads. A) Diagram of synaptic RGC->LIN->TC synaptic triad. Excitatory synapses (+), inhibitory synapses (-). B) EM showing RGC->LIN->TC triad. C) Distribution of the number of triadic relationships (RGC->TC + LIN1->TC pairs) within 5 μm of each of 135 analyzed RGC->LIN1 synapses. D) In 96 sampled RGC->LIN1->TC triads, the input to LIN1 usually occurs within a few microns of the output from LIN1.

To determine whether the proximity of the synaptic components of RGC->LIN->TC triad motifs reflected selective positioning of synapses or simply the density of synapses on LIN1 (a subcellular Peters Rule, (Peters and Feldman, 1976)), we measured RGC->LIN1 and LIN1->TC synapse proximity in random redistributions of TC synapses across LIN1. In the redistributions, only 57.2% of RGCs were within 5 μm of a TC synapse (95%CI 50.6-63.7%, compared to 84% observed, P < 0.001) and only 45.0% of TC synapses were within 5 μm of an RGC synapse (95%CI 40.5-49.6%, compared to 78% observed, P < 0.001). Thus, RGC inputs are distributed relative to TC outputs such that feedforward inhibition has a shorter path through the LIN1 arbor than predicted from a random distribution of synapses. Therefor independently overlapped distributions of RGC->LIN1 and LIN1->TC synapses are not sufficient to explain the tight associations of the components of triadic synapses

#### Lower frequency of RGC->LIN1->TC triad motif in the shell

The association of RGC and TC synapses with LIN was not evenly distributed across LIN1’s dendritic arbor. The density of non-triadic RGC inputs to LIN1 (i.e., that were not associated with nearby TC synapses) was significantly higher (3.61 times) in the dLGN shell (0.0296/μm, 32 synapses) compared to the density in the dLGN core (0.0082/μm, 39 synapses,) suggesting that LIN1 might interact differently in different parts of the dLGN (Figure 4A). Monte Carlo redistribution of the RGC-only motifs shows that this imbalance is unlikely to occur in a random distribution of independent synapses (mean ratio 1.02, 95%CI 0.48-1.73, P < 0.001), but could still be due to a difference in behavior of a subset or of different classes of RGCs in the shell.

#### LIN1->LIN->TC triad motif on the axon

The lack of RGC input on the axon-like neurite means that its profile of synaptic motifs will be distinct from that of the other neurites. The formation of LIN->TC synapses in the absence of RGC inputs (18/18 on the axon), only occurred in 71/724 TC outputs elsewhere on the arbor (ranksum P < 0.001). Interestingly, in the absence of RGC input, the LIN1 axon-like neurite often played an RGC-like role in LIN1->LIN->TC synaptic motifs (Figure 4I). LIN1 boutons of the axon formed nearby synapses onto both a TC and a presynaptic dendrite of another LIN. The postsynaptic LIN neurites also innervated the TC dendrite. The number of LIN1->LIN->TC motifs was much higher on the axon (16) than predicted by random redistributions of synapses across the LIN1 arbor (mean = 2.1, 95%CI 0-5, P <0.001). The function of such a disinhibitory triad is unclear, but the motif could play an analogous role to RGC triads, temporally shortening LIN1s inhibition to a subset of its target cells.

#### LIN->LIN motifs

Motifs between LINs included simple one-way inhibition, reciprocal synapses, autaptic synapses, and synaptic triads (Figure 2E, G Figure 4). The pre- and postsynaptic sites of the autaptic synapses were separated by <10 μm of arbor length and are therefore unlikely to constitute contacts between different functional domains of the same cell. We also checked whether the two sides of reciprocal and autaptic synapses were consistently innervated by either the same RGC bouton or innervated the same TC dendrite (reciprocal triads). While we observed both RGC->LIN<->LIN and LIN<->LIN->TC motifs associated with LIN1, we also observed reciprocal synapses with no nearby common partner. Thus, there is no clear indication that similarity of connectivity is required for the formation of reciprocal synapses.

#### Unidentified synapse motifs

Many of the non-RGC, non-LIN inputs to LIN1 also innervated a TC dendrite that was innervated by LIN1, thereby forming a potential BS->LIN->TC triadic motif (Figure 4K, described previously in cat (Sherman, 2004)). The one brainstem synapse that innervated the axon-like neurite of LIN1 was notable because that synaptic bouton innervated a neurite of another LIN that was also innervated by the LIN1 axon. Such a BS->LIN->LIN motif had been suggested by physiological evidence from cat dLGN (Cox and Sherman, 2000).

LIN1 therefore, engaged in many different synaptic relationships with many kinds of partners. LIN1 formed local triadic synaptic motifs with each type of input that formed synapses on its arbor. Variation in LIN->LIN connections included all the three-cell permutation of local connectivity pattern we could test for (RGC->LIN->LIN, LIN->LIN->TC, RGC->LIN<->LIN, LIN<->LIN->TC). While RGC inputs were consistent in forming triad synapses, there were still many TC outputs that were isolated from RGC inputs arguing for multiple modes of LIN->TC transmission.

### Neurite morphology drives distinct output modes in one LIN

Given the distinct morphologies of the trunk and targeting dendrites, we next asked whether these neurites played different roles in how they connect LINs to dLGN circuits.

#### Fasciculation with synaptic partners

Trunk dendrites formed synapses without any change in the path of the pre or postsynaptic neurite, and often without any morphological differentiation such as a bouton or spine (Figure 6A-B, Supplementary Movie 4). This lack of morphological differentiation suggests that the synapse formation on trunk dendrites is incidental, occurring when LINs happen to be adjacent to TC dendrites or RGC axons rather than by directed growth of the pre- or postsynaptic partners to a common site. In contrast, we found that the distinct circuitous paths of the targeting dendrites reflect fasciculation with synaptically connected RGC axons and TC dendrites (Figure 6C-D, Supplementary Movie 4,5). In this way, targeting dendrites brought LIN1 into extended contact with specific subsets of TC dendrites and bundles of RGC axons. To quantify fasciculation between trunk and targeting dendrites, we tested how far away LIN1 and an innervated TC stayed in proximity (<1μm) around the site of a synapse. We found that 12.0% (14/116) of synapses on trunk dendrites and 36.8% (56/152) on targeting dendrites maintained proximity for at least 5 μm (Figure 6E, Bootstrap difference in percentages = 0, 95%CI -11.1-10.2, P<0001). Therefore, synapses of targeting dendrites were associated with more opportunities for LIN1 to interact with the synaptic partner and the local microcircuitry. Of 60 targeting neurites tested, 32 included at least one synapse that met this criterion.

**Figure 6.**
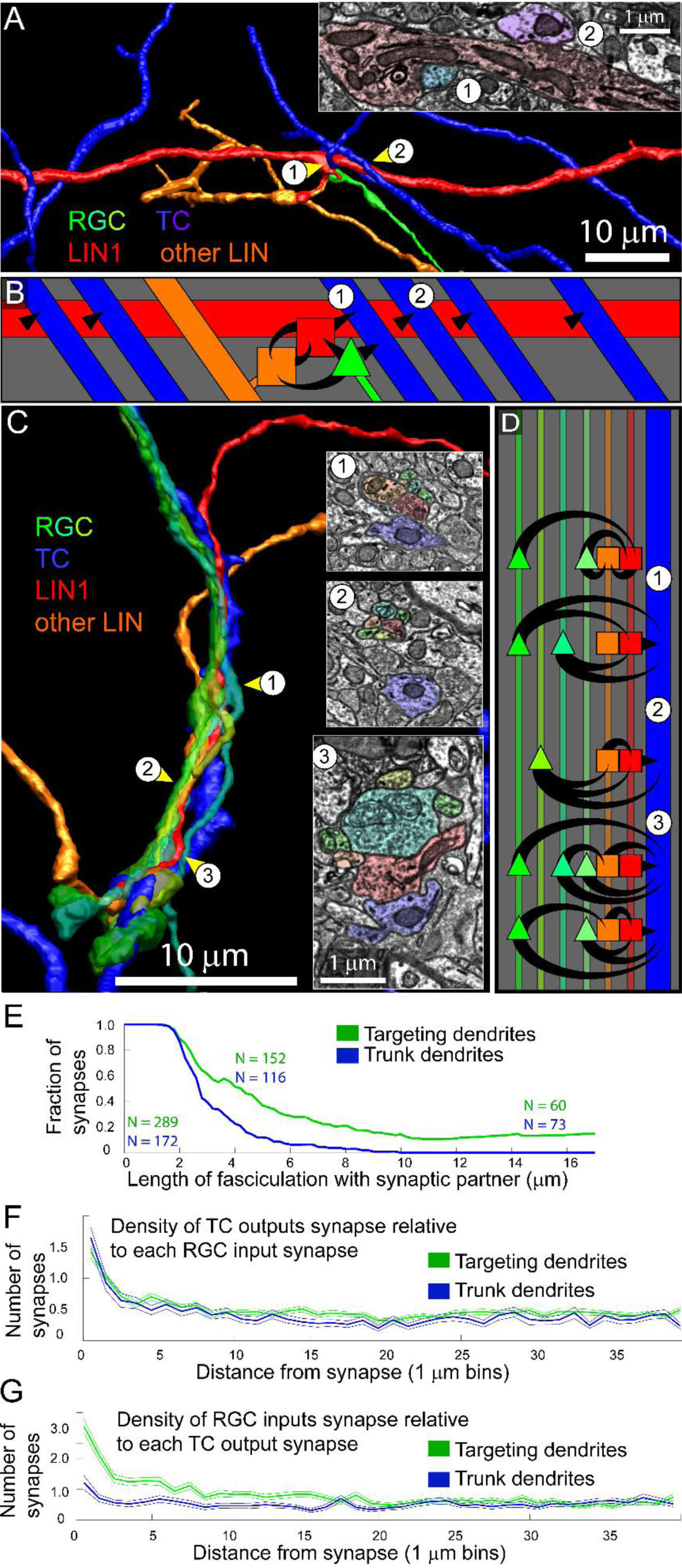
Distinct synaptic connectivities are generated by trunk and targeting dendrites of LIN1. A) Rendering of LIN1 trunk dendrite (red) and synaptically connected TCs (blue), LIN (orange), and RGCs (green). A short side branch of the LIN1 trunk dendrite receives input from the RGC and forms reciprocal synapses with a second LIN. Circled numbers indicate position of the two en-passant trunk synapses shown in the inset EM. B) Diagram of the simple/diffuse connectivity of the trunk dendrite in panel A. C) Rendering of a LIN1 targeting dendrite (red) and synaptically connected TC (blue), LIN (orange), and four RGC axons (green, lime, cyan, yellow). All six presynaptic neurites follow the TC dendrite through the neuropil. Circled numbers indicate position of the inset EM. D) Diagram of the complex/strong connectivity of the targeting dendrite in panel C. E) Length of fasciculation (LIN1 & TC <1μm away) relative to LIN1->TC synapses formed on trunk (blue) and targeting (green) dendrites. F) Change in density of LIN1->TC synapses with distance relative to RGC inputs onto trunk (blue) and targeting (green) dendrites. G) Change of density of RGC->LIN1 synapses with distance measured relative to LIN->TC synapses from trunk (blue) and targeting (green) dendrites.

#### Convergence

The fasciculated bundles of the LIN neurites, RGC axons, and TC dendrites were replete with synapses (Figure 6C, D) so that synaptic partners were often connected by multiple synapses and most neurites participated in multiple triadic motifs (Figure 6D). To quantify whether targeting dendrites of LIN1 were more likely to form multiple synapses on target TCs than trunk dendrites, we measured the rate at which nearby (≤10 μm) LIN1->TC synapses innervate the same TCs. We found that pairs of nearby LIN1->TC synapses from targeting dendrites innervated the same TC at a higher rate (33% of 318 synapse comparisons, 2.40 times higher) than pairs of synapses on trunk dendrites (14% of 232 synapse comparisons, bootstrap reassignment of trunk/targeting identity mean ratio = 1.01, 95%CI = 0.75-1.30, P<0.001). The increased probability of convergence onto target cells provides targeting dendrites with stronger (multisynaptic) connections to their synaptic partners than are found on trunk dendrites.

#### Motifs

To determine if the type of motifs generated by targeting and trunk dendrites differed, we compared the distances between potential triad input synapses (RGC->LIN synapses) and potential triad output synapses (LIN1->TC synapses). We found that trunk dendrites often formed synapses onto TCs in the absence of nearby RGC->LIN1 input (Figure 6F, G). Specifically, 28.2% (46/163 analyzed) of *trunk* LIN1->TC synapses were >5 μm away from RGC->LIN1 synapses, whereas only 4.5% (25/561 analyzed) of LIN1->TC synapses on *targeting* dendrites were isolated from RGC inputs. This difference (23.7%) was never observed in a Monte Carlo redistribution of synapses (mean = 0% difference, 95%CI = -5.0 to 5.0%, P < 0.001). Hence, the RGC input driving LIN->TC synapses of targeting dendrites is more local than that of trunk dendrites.

The above comparison shows, remarkably, that two types of dendrites of the same cell, embedded in the same neuropil, receiving and establishing the same types of synapses, with the same pre- and postsynaptic partners can, nonetheless, have quite different functions. The incidental en-passant relationships of trunk dendrites with their pre- and postsynaptic partners results in weak and solitary associations with whichever RGCs and TCs they pass by. In contrast, the fasciculation of targeting dendrites with their synaptic partners leads to strong multi-synaptic associations among specific cohorts of pre- and postsynaptic neurons. The extended structural proximity of multiple RGCs, LINs and TCs gives rise to complex chains of serial synapses and sites at which different sources of retinal input converge on the same local microcircuit. Given that most LIN->TC synapses occur near the TC cell body, the TC targets of these local microcircuits will be different from one region of LIN1’s arbor to another. The result of the two distinct dendrite types is that one LIN provides diffuse global inhibition to neurons in broad swaths of the dLGN while at the same time participating in highly targeted microcircuits for complex local computations.

## DISCUSSION

LINs in the dLGN have been studied for decades in a range of species including cats, primates and rodents (Bickford et al., 1999, 2010; Blitz and Regehr, 2005; Bloomfield and Sherman, 1989; Cox and Beatty, 2017; Crandall and Cox, 2012, 2013; Dankowski and Bickford, 2003; Halnes et al., 2011; Hámori et al., 1974; Hamos et al., 1985; Hirsch et al., 2015; Lam et al., 2005; Pasik et al., 1973, 1976; Seabrook et al., 2013; Wilson, 1989). Our connectivity map of a LIN in the mouse dLGN is broadly consistent with the conclusions reached in this body of work. What is new is that by doing circuit-scale serial electron microscopy we could assess almost all the pre- and postsynaptic interactions of an individual LIN. We found that the diverse range of functional attributes and synaptic relationships attributed to LINs in the visual thalamus as a class, remarkably, were present among the neurites of one single neuron. At a macroscopic level, the neuron we analyzed had synaptic associations across different functional and retinotopic domains. At the microscopic level, it utilized a diverse set of inhibitory synaptic motifs that included every type of local inhibition previously described in the visual thalamus. In sum, we find it difficult to assign a unitary role of this LIN because it participated in almost every conceivable kind of inhibition with every cell type and throughout many parts of the dLGN.

### Global promiscuity vs. local specificity

At the level of region-to-region connectivity, LIN1’s neurites followed Peter’s Rule (Peters and Feldman, 1976) in that the probability of synaptic connection within a region was reliably predicted by it’s arbor (Figure 3A). Saturated segmentation of the full volume would allow a more robust statistical test of this lack of specificity (Kasthuri et al., 2015; Mishchenko et al., 2010). At a finer scale, however, the morphological differentiation of targeting dendrites produced a much more specific connectivity pattern. The targeting between RGC, LIN and TC neurites produced extended bundles of functionally related neurites and the dense clusters of synapses characteristic of the classic EM profiles of dLGN synaptic glomeruli (Szentágothai, 1963). This morphological targeting concentrates the input and output synapses of triad synapses and clusters of triad synapses together on the scale of ∼2-20 μm consistent with the idea that individual neurites are forming discrete computational units in the dLGN. However, the importance of the distance between synapses depends entirely on how much signals attenuate across the arbor of LINs.

It is difficult to predict how far signals will spread in the targeting dendrites of LINs. Glutamate uncaging at distal LIN dendrites can drive inhibition of nearby TCs without depolarizing the LIN cell body (Crandall and Cox, 2012). Modeling of signal attenuation in LINs predicts that inputs to distal dendrites should be attenuated at the soma from ∼50% to 99% (Bloomfield et al., 1987; Briska et al., 2003; Halnes et al., 2011; Perreault and Raastad, 2006). The level of attenuation in these models varies depending on membrane resistance, process diameter, active conductances and the spatial and temporal pattern of the input. The varicose targeting dendrites, with their fine (< 200 nm) inter-varicosity diameter, may have more in common electrotonically with the varicose input/output neurites of AII amacrine neurons than to the trunk dendrites connecting distal neurites to the cell body of LINs. A study of these AII input/output neurites (Grimes et al., 2010) found that under naturalistic transient stimulation, the activity of other boutons was only relevant within about 20 μm, resulting in 150 parallel microcircuits in the same neuron. A similar length restriction in the targeting dendrites of LINs would mean that most of the inhibition provided by targeting LIN dendrites would be glomerulus specific.

Insight into the computational benefit of these functional units may be revealed by their synaptic organization. The fasciculation of a LIN1 targeting side branch with multiple RGC axons means that feedforward triadic motifs of many RGC axons pass through the same local patch of LIN1 neurite and TC dendrite. Individual RGC->LIN->TC triads are thought to narrow the window during which TCs can spike in response to RGC input (Blitz and Regehr, 2005). By extension, a complex of RGC->LIN->TC triads converging on the same TC might narrow the window during which overlapping RGC activity can sum to drive the TC to spike. Given that visual events tend to increase RGC synchronicity (Usrey and Reid, 1999), these dLGN microcircuits might be expected to selectively pass visual events over network noise. The potential for local microcircuits to selectively pass coincident spikes is made more interesting by the observation that nearby RGC boutons tend to share similar feature selectivities even though they are not always of the same RGC subtype (Liang et al., 2018). Therefore, the convergence of RGCs with diverse but overlapping receptive field properties in the same glomerulus should produce responses in TCs that are selective to the spatial and feature overlap of the converging RGCs. In this way, the multi-synaptic motifs observed in our serial electron microscopy data could generate receptive field properties that are more refined than those in the RGCs themselves.

### One neuron, many functions

The most remarkable thing about LIN1 was not the specific synaptic motifs we observed, but that so many distinct synaptic relationships were observed within a single neuron. Because locally integrated signals can be transmitted on a neurite-by-neurite basis in LIN1, this diversity in connections does not necessarily mean that LIN1 has a complex ensemble function, but rather that LIN1 performs many different functions. The combination of the capacity for local transmission of signals, the multi-region arborization, and the range of microcircuit motifs found along its arbor means it is difficult to count the number of visual computations this one neuron may participate in.

The emphasis on the cell-centric organization in the nervous system is one of the principle doctrinal ideas of Cajal (interestingly, strongly disputed by his co-Nobelist, Golgi). If function is divisible into local independent sites, then the fact that all these sites are associated with the same cell (i.e., supported by the same nucleus and cell body synthetic machinery) may be tangential to their function. In these cases, the neuron doctrine hides the actual computation complexity of the circuit. In transitioning from the analysis of region to region computations of Golgi type 1 cells (long axon) to the local computations of Golgi type 2 cells (short or no axon), a more neuropil-centric approach is required.

## ACKNOWLEDGEMENTS

Image alignment was performed by Art Wetzel at Pittsburgh Supercomputing (low res) and Adi Peleg and Shuohong Wang (high res). We gratefully acknowledge support from the NEI (1R21EEYE030623-01), NIH/NINDS (1DP2OD006514-01, TR01 1R01NS076467-01, 1U01NS090449-01, and U24NS109102), Conte (1P50MH094271-01), MURI Army Research Office (contract no. W911NF1210594 and IIS-1447786), NSF (OIA-1125087 and IIS-1110955), the Human Frontier Science Program (RGP0051/2014), the NIH and NIGMS via the National Center for Multiscale Modeling of Biological Systems (P41GM10371). Josh Morgan is a recipient of a Research to Prevent Blindness Career Development Award. This work was supported by an unrestricted grant to the Department of Ophthalmology and Visual Sciences from Research to Prevent Blindness and by National Eye Institute of the National Institutes of Health under award number P30 EY002687. We are grateful to James Cuff and Harvard Research Computing for data management. Thanks to Richard Schalek for microscope hardware management. Thanks to Katia Valkova for tracing help. Thanks to Jordan Matelsky, Brock Wester, William Gray-Roncal and the team at bossDB. Editing assistance povided by InPrint at Washington University in St. Louis.

## AUTHOR CONTRIBUTIONS

Conceptualization, J.L.M and J.W.L.; Methodology, J.L.M.; Software, J.L.M.; Formal Analysis, J.L.M, Investigation, J.L.M; Resources, J.L.M and J.W.L.; Data Curation, J.L.M.; Writing, J.L.M and J.W.L; Visualization, J.L.M.; Funding Acquisition, J.L.M and J.W.L;

## STAR METHODS

### Lead contact and materials availability

Further information and requests for resources and datasets should be directed to and will be fulfilled by the Lead Contact, Josh Morgan. This study did not generate unique reagents.

### Experimental model and subject details

All animals were handled according to protocols approved by the Institutional Animal Care and Use Committee at Harvard University. Tissue from the monocular zone of the dLGN was obtained from a 32 day old C57BL/6 mouse.

### Method details

The acquisition of the dLGN EM image volume used here was described in a previous publication (Morgan et al., 2016). Briefly, a P32 C57BL/6 mouse (gender unrecorded) was anesthetized with pentobarbital and perfused with 2% paraformaldehyde plus 2% glutaraldehyde. A 300 μm vibratome slice of dLGN was stained sequentially with 2% osmium tetroxide, followed by 0.1% thiocarbyhydrazide, then 2% osmium tetroxide (second layer binding to the first osmium-thiocarbyhydrazide complex, OTO, (Tapia et al., 2012), and 4% uranyl acetate. The vibratome section was dehydrated with acetonitrile, embedded in Epon-Araldite and sectioned at 30 nm using ATUM. Ten thousand sections were mounted on 4-inch silicon wafers (∼200 sections each) and were post-stained with 3% lead citrate. Images were acquired using the in-lens detector of a Zeiss Merlin SEM and our WaferMapper custom Matlab (Mathworks) code for automated imaging (Hayworth et al., 2014). The image stack was initially aligned by Art Wetzel at Pittsburgh Supercomputing (Wetzel et al., 2017). Images were acquired at 4 nm pixel size and downsampled to 16 nm pixel size for most of the image annotation. Image annotation (cell tracing, and synapse labeling) was performed manually using VAST (Berger et al., 2018) (https://software.rc.fas.harvard.edu/lichtman/vast/).

### Quantification and statistical analysis

Distances between synapses were calculated in two ways: Euclidian (3D) and topological (arbor based). Topological distances were calculated by first skeletonizing LIN1 using a shortest path algorithm described previously (Morgan et al., 2016). When searching LIN1 for synaptic motifs, we looked for groups of synapses in which the relevant LIN1 synapses were within 5 μm (topological distance) from one another. Relevant synapses not formed by LIN1 (as in the RGC->TC synapse of an RGC->LIN->TC triad) were included if they occurred within 5 μm (Euclidian distance) of the relevant LIN1 synapses. The tortuosity of each point of the LIN arbor was measured by identifying an approximately 5 μm length of neurite surrounding each node on the skeletonized LIN arbor. Tortuosity was defined as the total length of the neurite segment (∼ 5 μm) divided by the Euclidian distance between the endpoints of the neurite segment. Neurite diameter was measured by taking the smallest width of the 2D tracing surrounding each node of the skeletonized LIN arbor.

For the analysis of the probability of RGC input free neurites occurring, axon length stretches of the LIN1 arbor were automatically extracted by finding a LIN1 skeleton node furthest from the cell body and tracing back to include a 366 μm length of neurite (equal to the axon) connected to this endpoint. The procedure was then iterated on the next furthest untraced endpoint until no valid potential endpoint was present on the arbor.

For measures of fasciculation, pre and postsynaptic neurite proximity were measured relative to distance from synaptic contacts. To count the number of targeting neurites that exhibited fasciculation, the arbor was manually divided into branch segments longer than the scale of fasciculation analysis. Targeting dendrites were considered fasciculated if they stayed within 1 μm of its target TC dendrite 5 μm away from the site where they formed synapses.

### Data and code availability

All rendering and analysis were performed using custom Matlab code (Nautilus Analysis 4.0). This package of Matlab scripts imports segmentation files from VAST segmentation software and stores them as voxel lists. Cells and subcellular structures can then be retrieved individually for rendering or analysis. This code is available at https://sites.wustl.edu/morganlab/data-share/dlgn-2018/.

The dLGN dataset can be accessed at https://software.rc.fas.harvard.edu/lichtman/LGN/ and https://sites.wustl.edu/morganlab/data-share/dlgn-2018/. The segmentation files used in this publication can be accessed at https://sites.wustl.edu/morganlab/data-share/dlgn-2018/. The full resolution aligned dLGN dataset is available at http://bossdb.org/project/morgan2020.

## Additional resources

**See** https://sites.wustl.edu/morganlab/dlgn-lin1-resources/ for movies and additional resources

**Supplementary Movie 1**. LIN arbors, Related to Figure 1. Rotation of partial reconstructions of five LINs showing range of wide arbor orientations. Four of these LINs are shown in Figure 1B.

**Supplementary Movie 2**. LIN1, synapses, Related to Figure 2. Rotation of initially showing neurite types for LIN1, then input types, then output synapse types.

**Supplementary Movie 3**. 3D structure of LIN1 glomerulus. Related to Figure 4L. LIN1 is red. Within the glomerulus, TCs are blue, RGCs are green and other LINs are orange. Outside of the glomerulus, some tracings continue in grey. Scale bar is 10 μm.

**Supplementary Movie 4**. LIN1 trunk dendrite and synaptic partners, Related to Figure 6A. Rotation of trunk dendrite of LIN1 (red) and all its synaptic partners (RGC = green, TC = blue/purple, other LIN = orange) as also shown in Figure 6A. Cones point to position of synapses (RGC->LIN = yellow, LIN->TC = magenta, RGC->TC = cyan, LIN->LIN = red). Scale bar = 10 μm.

**Supplementary Movie 5**. LIN1 targeting dendrite and synaptic partners, Related to Figure 6C. Rotation of targeting presynaptic dendrite of LIN1 (red) and all of its synaptic partners (RGCs = green, TC = blue, other LIN = orange) as also shown in Figure 6B. Cones point to position of synapses (RGC->LIN = yellow, LIN->TC = magenta, RGC->TC = cyan, LIN->LIN = red). Scale bar = 10 μm.

## SUPPLEMENTAL INFORMATION

**Supplementary Figure 1.**
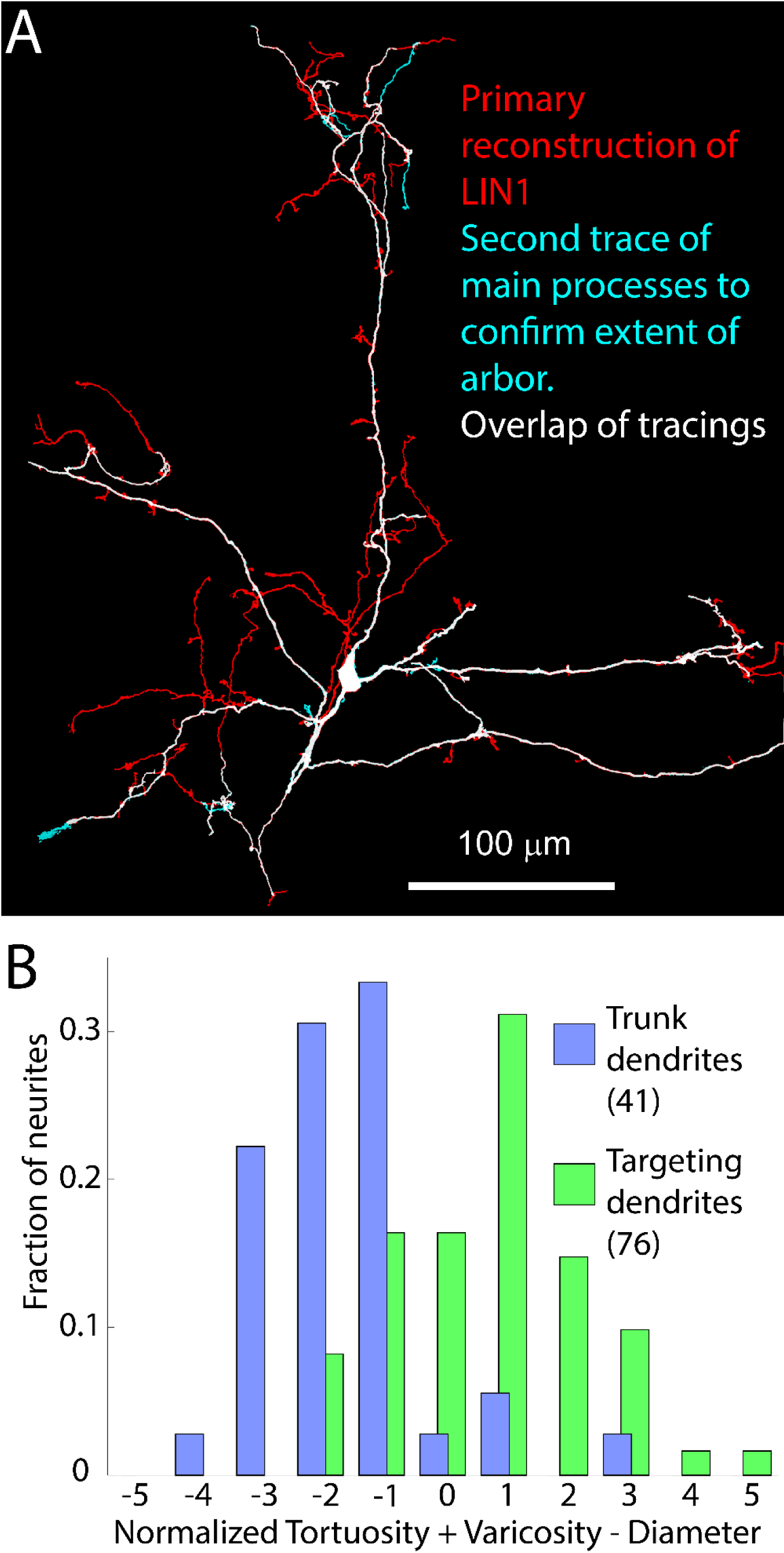
Morphology of LIN1, Related to Figure 1 A) An independent tracing (cyan) of the main processes of LIN1 was used to confirm the wide arborization observed in the original tracing (red). B) Morphology of neurites identified as trunk or targeting dendrites shown as combined measure of normalized ([x-mean]/standard deviation) tortuosity + varicosity – diameter.

**Supplementary Figure 2.**
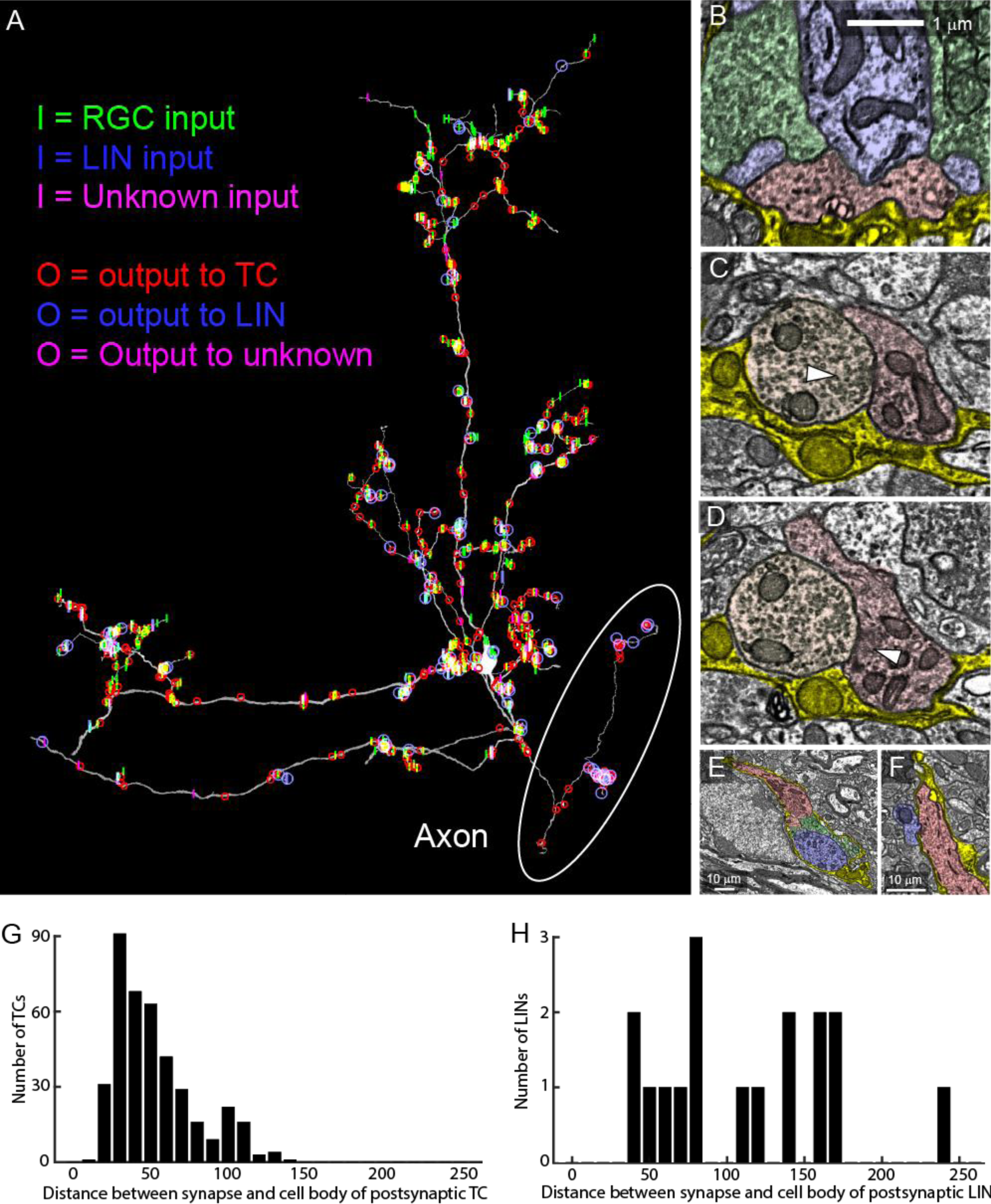
Synaptic connections of LIN1, Related to Figure 2. A) Distribution of different synapse types across LIN1. LIN1 is rotated to a sagittal view so that the axon is clearly distinguishable. Axon-like neurite is circled. B) Electron micrograph of a synapse formed by the axon of LIN1 (red) onto a TC dendrite (blue). RGCs that innervate spines of the TC are shown in green. Glial sheath is shown in yellow. (C-D) Reciprocal motif composed of a synapse from the axon of LIN1 onto another LIN (orange, C) and a synapse from another LIN onto LIN1 (D). E) Electron micrograph showing glial sheath of LIN1 is continuous with the glial sheath of a synaptic glomerulus innervated by LIN1. LIN1 (red), a second LIN (orange), RGCs (green), TC dendrite (blue) and glial ensheathment (yellow) are highlighted. F) Synaptic connection formed between trunk dendrite of LIN1 (red) and dendrite of TC (blue). The synapse is formed at a small gap in the glial ensheathment (yellow). G) Histogram of the distances between synapses between LIN1 and TCs and the cell body of the postsynaptic TC. Most synapses onto TCs were formed within 50 μm of the TC cell body. H) Histogram of distances between LIN1->LIN synapses and the cell bodies of LINs that could be traced to those synapses. Most LIN1->LIN synapses led to neurites that exit the EM volume.

**Supplemental Figure 3.**
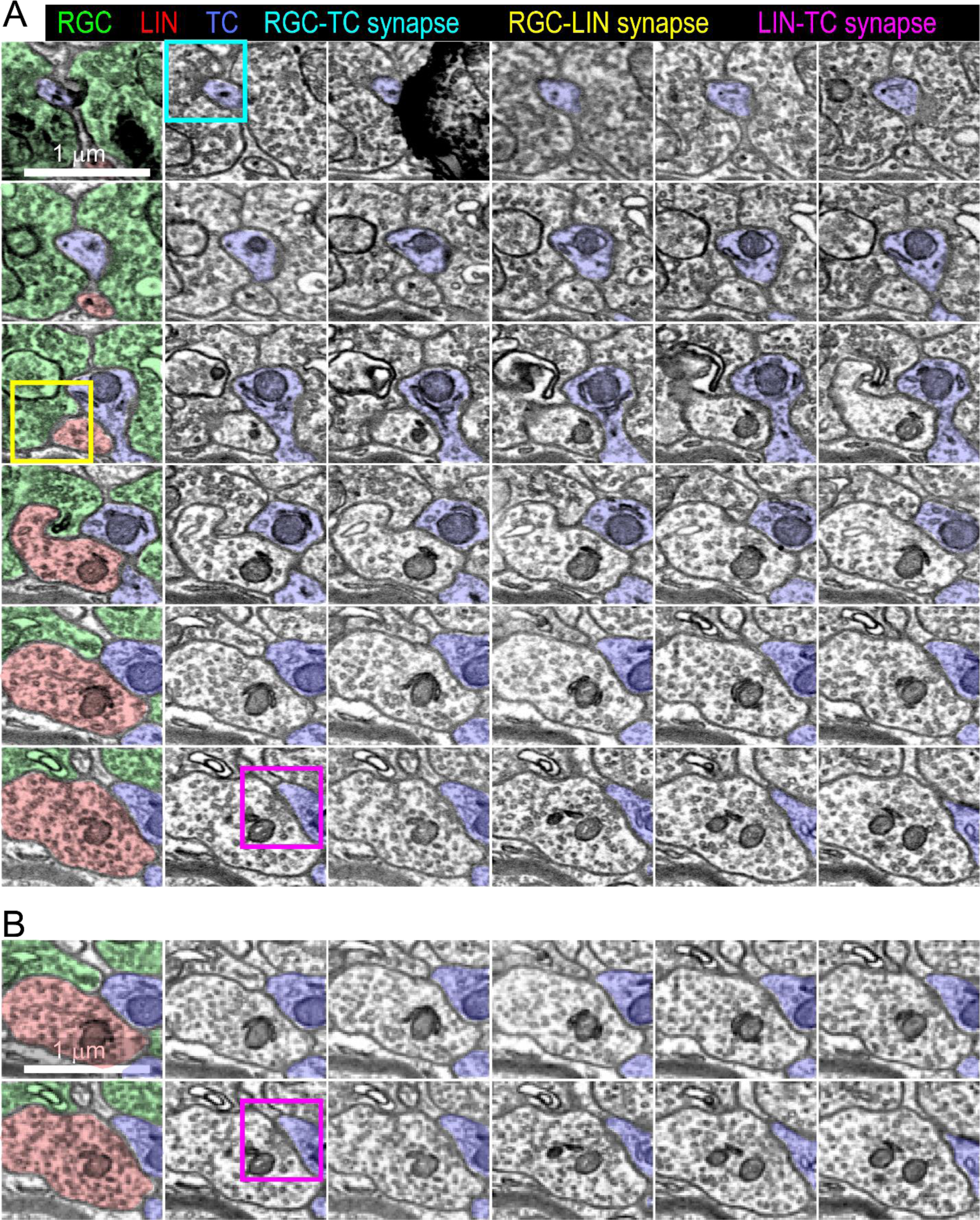
EM of a triad synapse, Related to Figure 3. Example of a synaptic triad and comparison between high resolution and downsampled images series through a synapse. A) EM series (full 4 nm x 4nm x 30 nm resolution) through an RGC->LIN->TC triadic synaptic arrangement. The postsynaptic TC is shown in blue. Cyan box = RGC->TC synapse. Yellow box = RGC->LIN synapse. Magenta box = LIN->TC synapse. B) Downsampled (16 nm x 16 nm x 30 nm) version of LIN1->TC synapse shown in A.

